# Impact of age-specific immunity on the timing and burden of the next Zika virus outbreak

**DOI:** 10.1101/661223

**Authors:** Michel J. Counotte, Christian L. Althaus, Nicola Low, Julien Riou

## Abstract

The 2015–2017 epidemics of Zika virus (ZIKV) in the Americas caused widespread protective immunity. The timing and burden of the next Zika virus outbreak remains unclear. We used an agent-based model to simulate the dynamics of age-specific immunity to ZIKV, and predict the future age-specific risk using data from Managua, Nicaragua. We also investigated the potential impact of a ZIKV vaccine. Assuming lifelong immunity, the risk of a ZIKV outbreak will remain low until 2035 and rise above 50% in 2047. The imbalance in age-specific immunity implies that people in the 15–29 age range will be at highest risk of infection during the next ZIKV outbreak, increasing the expected number of congenital abnormalities. ZIKV vaccine development and licensure are urgent to attain the maximum benefit in reducing the population-level risk of infection and the risk of adverse congenital outcomes. This urgency increases if immunity is not lifelong.

## 1 Introduction

Zika virus (ZIKV) is a flavivirus, which is transmitted primarily by mosquitoes of the genus *Aedes.* Before 2007, circulation of the virus only occurred sporadically in African and Asian countries (Wikan and Smith, 2016; Kohl and Gatherer, 2015). Between 2007 and 2013, ZIKV caused large-scale epidemics in the populations of Micronesia (Duffy et al., 2009), French Polynesia (Cao-Lormeau et al., 2014) and other Pacific islands (Wikan and Smith, 2016). ZIKV probably became established in *Aedes aegypti* mosquitoes in the Americas between 2013-2014, (Faria et al., 2016; Zhang et al., 2017) and then spread rapidly across the continent. In 2015, doctors in Brazil started reporting clusters of infants born with microcephaly, a severe congenital abnormality, and of adults with Guillain-Barré syndrome, a paralyzing neurological condition, resulting in the declaration by the World Health Organization (WHO) of a Public Health Emergency of International Concern (PHEIC) (World Health Organization, 2016). WHO stated, in September 2016, that ZIKV in pregnancy was the most likely cause of the clusters of microcephaly, and other adverse congenital outcomes (Krauer et al., 2017; Counotte et al., 2018). The risk of an affected pregnancy appears highest during the first trimester, with estimates between 1.0 and 4.5% (Cauchemez et al., 2016; Johansson et al., 2016). By the beginning of 2018, over 220,000 confirmed cases of ZIKV infection had been reported from Latin America and the Caribbean (PAHO, 2019), which is estimated to be only 1.02% (± 0.93%) of the total number of cases, based on mathematical modelling studies (Zhang et al., 2017).

Protective immunity conferred by infection, combined with high attack rates and herd immunity, can explain the ending of epidemics and the lack of early recurrence (Dietz, 1975), as has been seen with ZIKV (Ferguson et al., 2016). The duration of protective immunity induced by ZIKV infection remains uncertain, since immunity to ZIKV infection was not studied extensively before the 2013 outbreaks. Evidence from seroprevalence studies in French Polynesia and Fiji found that levels of ZIKV neutralizing antibodies decrease with time (Henderson et al., 2019). If the fall in antibody levels means that people become susceptible to infection again, population level ZIKV immunity might be declining already. Even if protective immunity is lifelong, the risk of a new ZIKV outbreak will rise as susceptible newborns replace older individuals, lowering the overall proportion of the population that is immune. A modelling study, based on data from the 2013 epidemic in French Polynesia, estimated that ZIKV outbreaks are unlikely to occur for 12 to 20 years, assuming lifelong immunity (Kucharski et al., 2016).

A direct consequence of population renewal will be an unequal distribution of immunity by age group, with younger age groups at higher risk from a new epidemic than older people (Ferguson et al., 2016). That effect will be amplified if ZIKV attack rates are lower in children than adults. Assessing the risk of ZIKV infection in women of reproductive age is essential because ZIKV infection in pregnancy, leading to adverse congenital outcomes, has such important implications for individuals, for public health and for investment in surveillance and mitigation strategies, including vector control, early warning systems, and vaccines (Abbink et al., 2018; World Health Organization, 2018). However, currently no vaccine is available against ZIKV. Phase I clinical trials of ZIKV candidate vaccines have shown levels of neutralizing antibody titers that were considered protective against reinfection (Gaudinski et al., 2018; Modjarrad et al., 2018). Some vaccines have already entered phase II trials (National Institute of Allergy and Infectious Diseases, 2018), but some companies have stopped vaccine development (Cohen, 2018).

Researchers in Managua, Nicaragua were the first to report the age-stratified seroprevalence of ZIKV antibodies in population-based surveys (Zambrana et al., 2018). The first cases of autochthonous ZIKV infection in Nicaragua were reported in January, 2016, and an epidemic was observed between July and December of that year. Through case-based surveillance, the public health authorities of Nicaragua reported a total of 2,795 people with ZIKV detected by reverse transcriptase (RT) PCR over this period (PAHO, 2019). The number of symptomatic infections is likely much higher, owing to under-reporting. Furthermore, ZIKV infection is asymptomatic in 33 to 87% of cases [23], which are generally not identified by surveillance systems. Shortly after the end of the 2016 epidemic, Zambrana et al. analyzed sera from two large population-based surveys in Managua to measure the prevalence of IgG antibodies against ZIKV in 2-to 14-year olds (N=3,740) and 15- to 80-year olds (N=2,147) (Zambrana et al., 2018). The authors reported ZIKV seroprevalence of 36.1% (95% confidence interval, CI: 34.5; 37.8%) among the 2-14 year age group and 56.4% (95% CI: 53.1; 59.6%) among the 15-80 year age group (Zambrana et al., 2018; Balmaseda et al., 2017). The observed post-outbreak seroprevalence in adults is in line with findings from seroprevalence studies from French Polynesia, Brazil, and Bolivia (Aubry et al., 2017; Netto et al., 2017; Saba Villarroel et al., 2018).

In this study, we used data from the 2016 ZIKV epidemic in Managua and developed an agent-based model (ABM) to predict the evolution of age-specific protective immunity to ZIKV infection in the population of Managua, Nicaragua during the period 2017–2097. We assessed: 1) the risk of a future ZIKV outbreak; 2) the consequences of a future ZIKV outbreak on women of reproductive age; 3) the influence of loss of immunity on future attack rates; and 4) how vaccination could prevent future ZIKV outbreaks.

## 2 Methods

### 2.1 Modelling strategy

We assessed the consequences of future outbreaks of ZIKV infection in Managua, Nicaragua using a stochastic ABM. The model follows a basic susceptible-infected-recovered (SIR) framework and integrates processes related to ZIKV transmission, immunity, demography, adverse congenital outcomes and vaccination (Table 1). We parameterized the model based on published estimates or inferences from data about the 2016 ZIKV epidemic (Table 1). We considered different scenarios about the duration of immunity, the timing and scale of ZIKV reintroductions in the population, and the timing and scale of a hypothetical vaccination program targeted towards 15 year old girls.

**Table 1:** Parametrization of the agent-based model. *^a^*age-dependent parameters; *^b^*the different scenarios are discussed in the text in detail under the headings corresponding to the headings of this table.

| Parameter | Comment | Source |
| --- | --- | --- |
| **ZIKV epidemic parameters** |  |  |
| Transmission rate<sup>a</sup> | Inferred from the 2016 epidemic | Zambrana et al. (2018) |
| Recovery rate | Inferred from the 2016 epidemic | Zambrana et al. (2018) |
| **ZIKV immunity** |  |  |
| Initial immunity<sup>a</sup> | Inferred from the 2016 epidemic | Zambrana et al. (2018) |
| Duration of immunity | Lifelong or decaying with time | 5 scenarios<sup>b</sup> |
| **Demography** |  |  |
| Initial age distribution | – | World Bank (2019a) |
| Birth rate | – | World Bank (2019a) |
| Death rate<sup>a</sup> | – | World Health Organization (2019) |
| Ageing | Linear ageing at each time-step | – |
| **ZIKV reintroduction** |  |  |
| Delay until reintroduction | 1 to 80 years | 80 scenarios<sup>b</sup> |
| Cases reintroduced | 1, 5 or 10 cases | 3 scenarios<sup>b</sup> |
| **Risk of adverse congenital event** |  |  |
| Exposure | Proportion of women in the first semester of pregnancy | World Bank (2019a) |
| Risk of microcephaly | Upon infection during exposure time (3 levels of risk) | Cauchemez et al. (2016); Johansson et al. (2016) |
| **Targeted vaccination** |  |  |
| Date of implementation | In 2021, 2025 or 2031 | 3 scenarios<sup>b</sup> |
| Effective coverage | Proportion of 15 year old girls vaccinated (0% to 80%) | 5 scenarios<sup>b</sup> |

### 2.2 Model structure

We simulated a population of 10,000 individuals for 80 years (2017–2097). We assigned agents age and ZIKV infection status (susceptible *S*, infected *I* or immune R). Initial conditions reflected the situation in Managua, Nicaragua in 2017, when there was no documentation of active transmission. In the outbreak-free period, we only considered demographic and immunity processes: births, deaths, ageing and, if applicable, loss of immunity and vaccination. Given the scarcity of these events at the individual level, we select a long time-step of seven days and stochastically applied the transition probabilities at each time step for each agent. After a given time, ZIKV-infected cases were reintroduced in the population. Upon reintroduction, the time step was reduced to 0.1 days, and we evaluated the epidemic-related transition probabilities: Susceptible agents may become infected at a rate *β_a_I/N*, where *β_a_* is the age-dependent transmission rate and N the total population size. Infected individuals may recover with a rate 7. We ignored the influence of the vector population and assumed that the force of infection is directly proportional to the overall proportion of infected individuals. We allowed six months for the outbreak to finish after introduction. Simulations were conducted independently for each combination of scenarios and repeated 1,000 times. In the baseline scenario, we assumed no vaccination, no loss of immunity and a reintroduction of 10 infected individuals.

We implemented the model in ‘Stan’ version 2.18 (Carpenter et al., 2017) and we conducted analyses with R version 3.5.1 (R Core Team and Team, 2008). The Bayesian inference framework Stan permits the use of probability distributions over parameters instead of single values, allowing for the direct propagation of uncertainty. Stan models are compiled in C++, which improves the efficiency of simulations. Algorithm 1 (Appendix A.1) describes the ABM in pseudo code. The model code and data are available from http://github.com/ZikaProject/SeroProject.

### 2.3 Parametrization

#### 2.3.1 ZIKV epidemic parameters

We inferred the probability distributions for the age-specific transmission rate *β_a_* and the recovery rate γ from data on the 2016 ZIKV epidemic in Managua, Nicaragua. We used surveillance data (Zambrana et al., 2018), which give weekly numbers of incident ZIKV infections, confirmed by RT-PCR (dataset 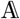, n=1,165), and survey data on age-stratified ZIKV seroprevalence, measured among participants of pediatric and household cohort studies in Managua during weeks 5–32 of 2017 (dataset 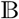, n=3,740 children and 1,074 adults) (Zambrana et al., 2018).

We conducted statistical inference using a deterministic, ordinary differential equation (ODE)-based version of the ABM with three compartments (*S, I* and *R*) and two age classes (*a* ∈ {1, 2} corresponding to ages 0–14 and ≥15):

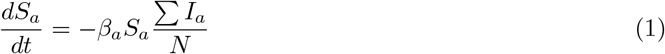

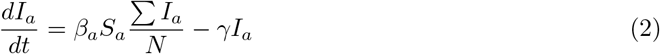

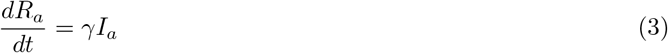

We ignored demography in this model because it covers a short time span. We recorded the overall cumulative incidence of ZIKV cases using a dummy compartment:

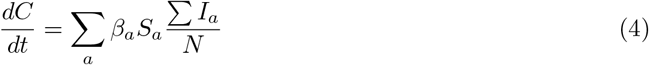

in order to compute the weekly incidence on week *t*:

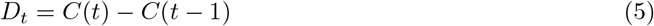

We fitted the model to weekly incidence data 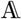 using a normal likelihood after a square-root variance-stabilizing transformation (Guan, 2009):

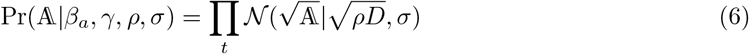

where *ρ* is a reporting rate parameter and *σ* an error parameter. In addition, we also fitted the model to the number of individuals with anti-ZIKV antibodies at the end of the epidemic by age group 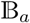 using a binomial likelihood:

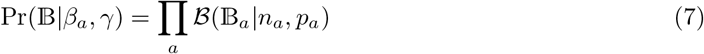

where 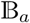 the number of individuals with antibodies, *n_a_* is the sample size in each age group, and *P_a_* = *R_a_*(*t_end_*)/*N_a_*(*t_end_*) the proportion of immune at the end of the epidemic. The full likelihood was obtained by multiplying Eq. 6 and Eq. 7. We chose weakly-informative priors for all parameters and fitted the model in Stan (Table 2). We describe the calculation of the basic reproduction number R_0_ in appendix A.2. We used one thousand posterior samples for *β_α_* and *γ* obtained by Hamiltonian Monte Carlo in the ABM model, ensuring the full propagation of uncertainty. Parameter values can translate from deterministic to agent-based versions of an epidemic model if the time step is small (Roche et al., 2011a), which was the reason for using a time step of 0.1 days.

**Table 2:** Parameter estimates inferred from incidence and sero-prevalence data on the 2016 ZIKV epidemic in Managua, Nicaragua. CrI: Credible interval.

| Parameter | Interpretation | Prior | Posterior<br>(median and 95%<br>CrI) |
| --- | --- | --- | --- |
| $\beta_1$ | Transmission for age group 0-14 | Expon(0.1) | 0.19 (0.16; 0.22) |
| $\beta_2$ | Transmission for age group $\geq 15$ | Expon(0.1) | 0.32 (0.30; 0.36) |
| $1/\gamma$ | Duration of infectious period | Gamma(1, 0.1) | 4.8 (4.3; 5.4) |
| $\rho$ | Reporting rate | Beta(1, 1) | 0.24% (0.21; 0.26) |
| $I(0)$ | Initial number of infectious | Expon(0.1) | 74 (40; 134) |
| $\mathcal{R}_0$ | Basic reproduction number | – | 1.58 (1.56; 1.59) |

#### 2.3.2 ZIKV immunity

We used the deterministic model, described in the previous section, to infer the proportion of people with protective immunity within each age group at the end of the 2016 epidemic 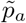. We used one thousand posterior samples of 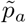 in the ABM to allow the propagation of uncertainty. Protective immunity to ZIKV after infection was lifelong in our first scenario, so the reduction of the overall proportion of immune individuals in the population decreased only because of population renewal. Given the absence of evidence about the duration of immunity to ZIKV, we considered four scenarios assuming exponentially distributed durations of immunity with means of 30, 60, 90, or 150 years. These values correspond to a proportion of initially immune agents that loses immunity after 10 years of 28%, 15%, 11% or 6%, respectively (Appendix A.3).

#### 2.3.3 Demography

We based the initial age distribution of the population on data from the World Bank (World Bank, 2019b). We used age-dependent death rates for 2016 from the World Health Organization (World Health Organization, 2019). For births, we computed a rate based on an average birth rate in Nicaragua of 2.2 births per woman, which was uniformly distributed over the female reproductive lifespan (World Bank, 2019a). We defined the period of reproductive age between 15 and 49 years. The ageing process was linear, increasing the age of each agent by 7 days at each 7-day time step.

#### 2.3.4 ZIKV reintroduction

We reintroduced ZIKV in the population after a delay of *d* = {1, …, 80} years in independent simulations. We chose this approach rather than continuous reintroductions to remove some of the stochasticity and assess more clearly the association between immunity decay and risk of an outbreak. As the probability of an extinction of the outbreak depends on the number of ZIKV cases reintroduced in the population, we considered three different values for the seed (1, 5 or 10 cases) and compared the results (Appendix A.4). Simulations using continuous reintroductions each year are presented in the appendix A.5.

#### 2.3.5 Risk of adverse congenital outcomes

The estimated number of microcephaly cases resulting from the reintroduction of ZIKV depended on the exposure, i.e. the number of pregnant women infected by ZIKV during their first trimester, to which we applied three different levels of risk, based on published estimates (Cauchemez et al., 2016; Johansson et al., 2016). We obtained the number of ZIKV infections among women aged 15–49 years from ABM simulations. As gender was not explicitly considered in the model, we assumed that women represented 50% of the population. We assumed a uniform distribution of births during the reproductive period, and considered that the first trimester constituted a third of ongoing pregnancies at a given time. We explored three different levels of risk of microcephaly in births to pregnant woman with ZIKV infection during the first trimester, as reported by Zhang et al., based on data from French Polynesia (0.95%, called low risk) and Brazil (2.19% and 4.52%, called intermediate and high risk, respectively) (Cauchemez et al., 2016; Johansson et al., 2016).

#### 2.3.6 Vaccination

We examined the effects of a potential ZIKV vaccine, given to 15-year-old-girls. This vaccination strategy was used for rubella virus, which also causes congenital abnormalities, before the vaccine was included in the measles, mumps and rubella vaccine given in childhood (Vyse et al., 2002). The main objective of vaccination would be the prevention of adverse congenital outcomes, including microcephaly. We simulated this intervention in the ABM, assuming vaccine implementation starting in 2021, 2025 or 2031. From that date, half of the agents reaching age 15, representing females, could transition to immune status *R* regardless of their initial status, with an effective vaccination coverage ranging from 20% to 80%.

### 2.4 Outcome analysis

From the simulations, we collected 1) the evolution of the age-specific ZIKV immunity in the population; 2) the attack rate resulting from the reintroduction of ZIKV at year d; 3) the age of newly infected individuals. We fitted a binary Gaussian mixture model to dichotomize the observed attack rates into either outbreaks or non-outbreaks. We defined the outbreak threshold as the 97.5% upper bound of the lower distribution. This corresponded to a threshold of 1%, so that attack rates ≥1% were considered as outbreaks. The age structure of newly infected individuals was used to compute relative risks of infection by age group.

## 3 Results

### 3.1 2016 ZIKV epidemic

The fitted model (Figure 1), resulted in a reporting rate of 0.24% (95% credible interval, CrI: 0.21; 0.26). The transmission rate in the 0–14 age group was 42% (95% CrI: 35; 48) lower than in the ≥1515 age group. This corresponded to an overall basic reproduction number of 1.58 (95% CrI: 1.56; 1.59). The predicted percentage of immune at the end of the epidemic was 36% (95% CrI: 34; 38) for the 0–14 age group and 53% (95% CrI: 50; 57) for the ≥15 age group.

**Figure 1:**
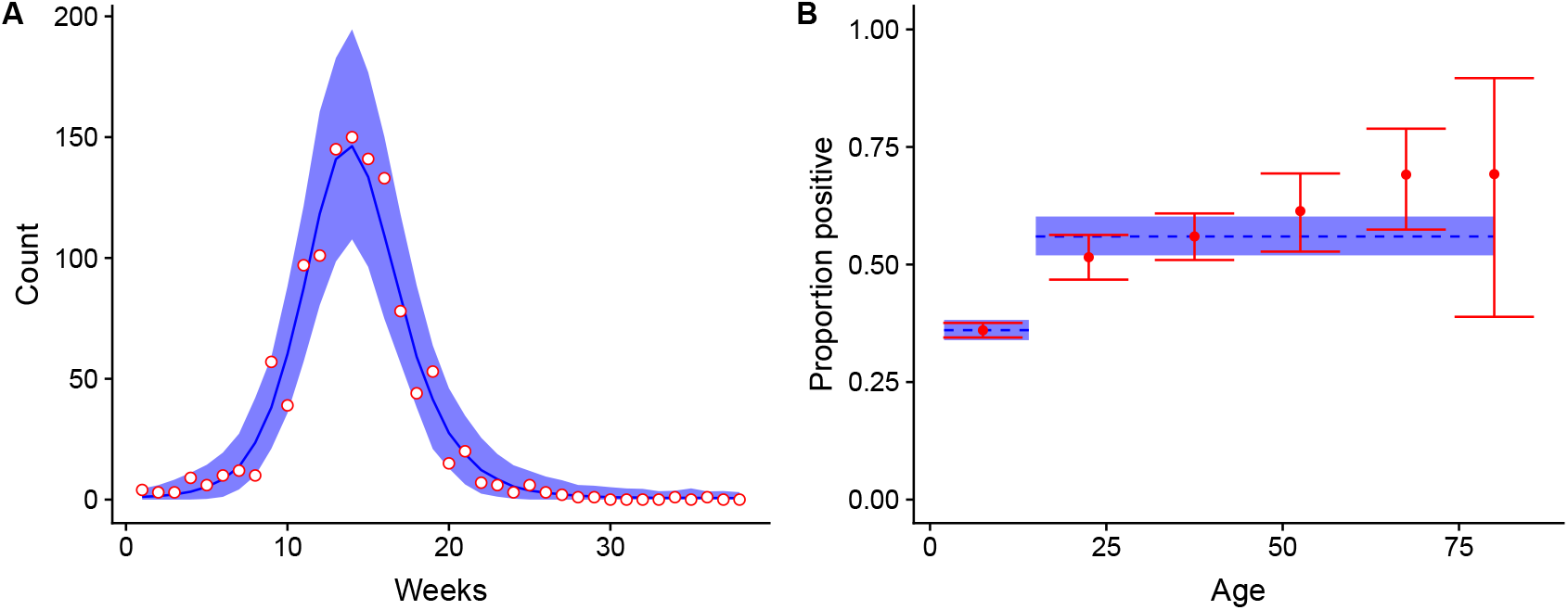
Model fit for (A) weekly incidence data and (B) post-epidemic sero-prevalence data from the 2016 ZIKV epidemic in Managua, Nicaragua. Data points are in red and the corresponding model fit (posterior median and 95% credible interval) is in blue.

### 3.2 Immunity and population

In our forward simulations, the expected population size increased by 42% between 2017 and 2097. Under the assumption that ZIKV infection results in lifelong protective immunity, population renewal will create an imbalance in the proportion immune in different age groups. We expect the overall proportion of the population with protective immunity to have halved (from 48% to 24%) by 2051 and to be concentrated among the older age classes (Fig. 2A). The 0–14 year old age group will become entirely susceptible by 2031 and the 15-29 year old age group by 2046.

### 3.3 Future risk of ZIKV outbreak

Reintroductions of ZIKV in the population of Managua are unlikely to develop into sizeable outbreaks before 2035, 24 years after the 2016 epidemic, assuming lifelong immunity for individuals infected in 2016 (Fig. 2B). After this point, attack rates resulting from ZIKV reintroduction will rise steeply. By 2047, we predict that ZIKV reintroductions will have a 50% probability of resulting in outbreaks with attack rates greater than 1% (Fig. 2C).

**Figure 2:**
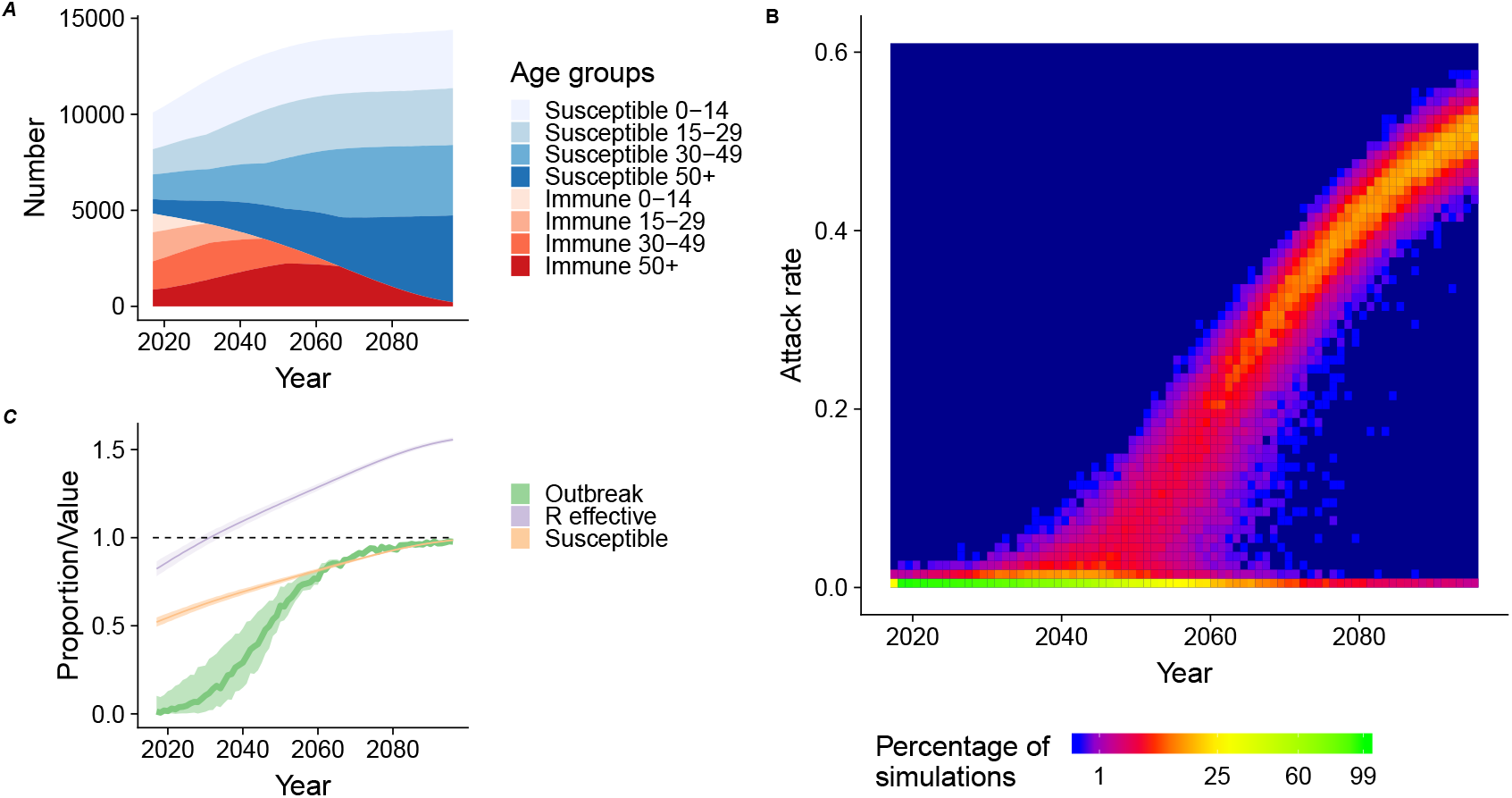
(A) The evolution of the immunity status per age group in a population of 10,000 agents for the next 80 years based on the demographic structure of Nicaragua. (B) Heat map of the distribution of the attack rates resulting from the reintroduction of ZIKV in the population at each year (1000 simulations for each year). (C) The evolution of the proportion of reintroductions resulting in outbreaks (with a threshold of 1%) with time (green), proportion of susceptible (orange), and effective reproduction number 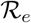 (purple).

### 3.4 Risk of infection and microcephaly births in women of reproductive age

The differences between age groups in both immunity and transmission will result in a disproportionate burden of infection in the 15–29 age class. The relative risk of infection in this age group ranges from 1.2 to 1.6, compared with the general population if an outbreak occurs during the period 2032–2075 (Fig. 3A). As most pregnancies occur in this age group, these women are also the most likely to experience a pregnancy with an adverse outcome. The increased risk of infection in this group implies that the number of adverse congenital outcomes resulting from a ZIKV outbreak during this period is likely to be higher than expected with a homogeneous distribution of immunity across ages. Assuming different values for the added risk of microcephaly after a ZIKV infection during the first trimester, we expect the mean number of additional microcephaly cases due to ZIKV infection resulting from the reintroduction of the virus in Managua, Nicaragua to reach 1 to 5 cases per 100,000 population in 2060 (Fig. 3B).

**Figure 3:**
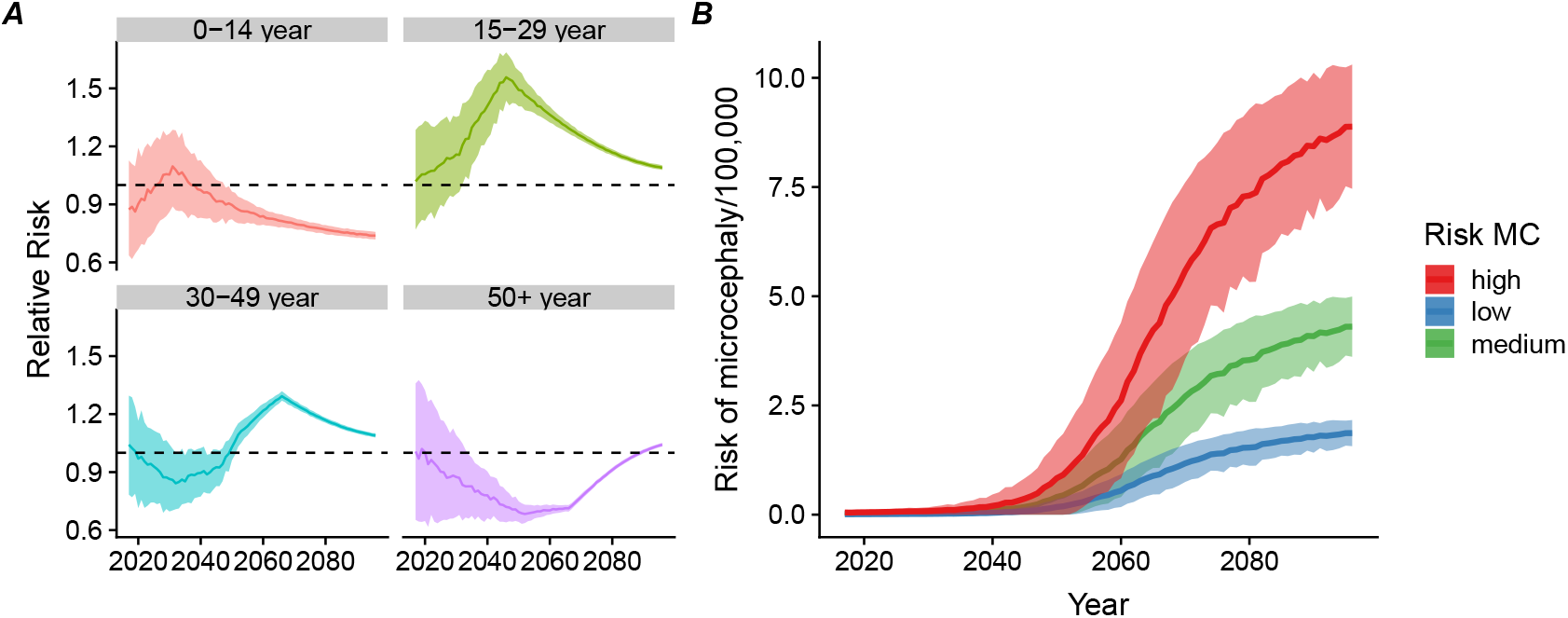
(A) Relative risk of ZIKV infection during a ZIKV outbreak per age group compared to the general population by year (median, interquartile range). (B) Expected number of additional microcephaly events associated with ZIKV infection during pregnancy per 100,000 total population according to three different risk scenarios.

### 3.5 Loss of immunity

If protective immunity to ZIKV is not lifelong, the time window before observing a rise in the attack rates resulting from ZIKV reintroduction will shorten (Fig. 4A). For instance, if 15% of the those who were infected in 2016 lose their immunity after 10 years (a mean duration of immunity of 60 years), the time until the risk of outbreak upon reintroduction reaches 50% would be 14 years earlier (2033) than with lifelong immunity (2047). Loss of immunity over time would reduce the relative risk in the 15–29 year old age group (Fig. 4B).

**Figure 4:**
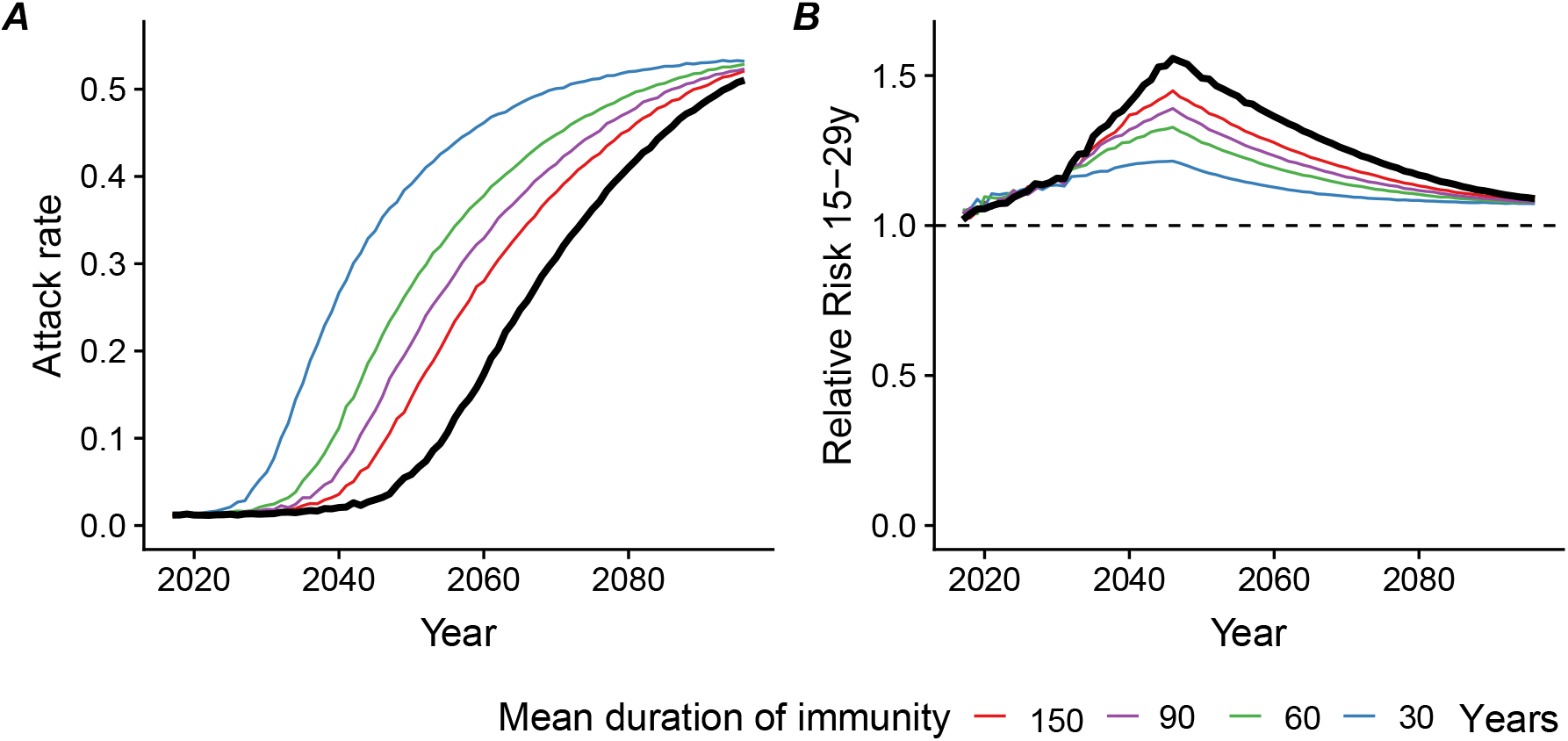
Consequences of alternative scenarios regarding the mean duration of protective immunity (30, 60 and 150 years), compared with lifelong immunity (thick black line): (A) median attack rate of ZIKV among reintroductions resulting in outbreaks (with a threshold of 1%) and (B) relative risk of ZIKV infection during an outbreak in the 15–29 year age group compared with the general population.

### 3.6 Targeted vaccination

The implementation of a vaccination program targeted towards 15 year old girls between 2021 and 2031 would reduce the risk of infection in women aged 15-29 years and would also indirectly reduce the overall risk of a ZIKV outbreak in the population (Fig. 5). If effective vaccine coverage is 60–80% amongst 15 year old girls, the prolongation of herd immunity could effectively mitigate the overall risk of a ZIKV outbreak in the population. The reduction in the number of microcephaly cases would then exceed what would be expected by considering only the direct protection granted by a vaccine to future mothers. A later implementation of the intervention would be less effective, as it becomes more difficult to maintain the herd immunity (Fig. 5B).

**Figure 5:**
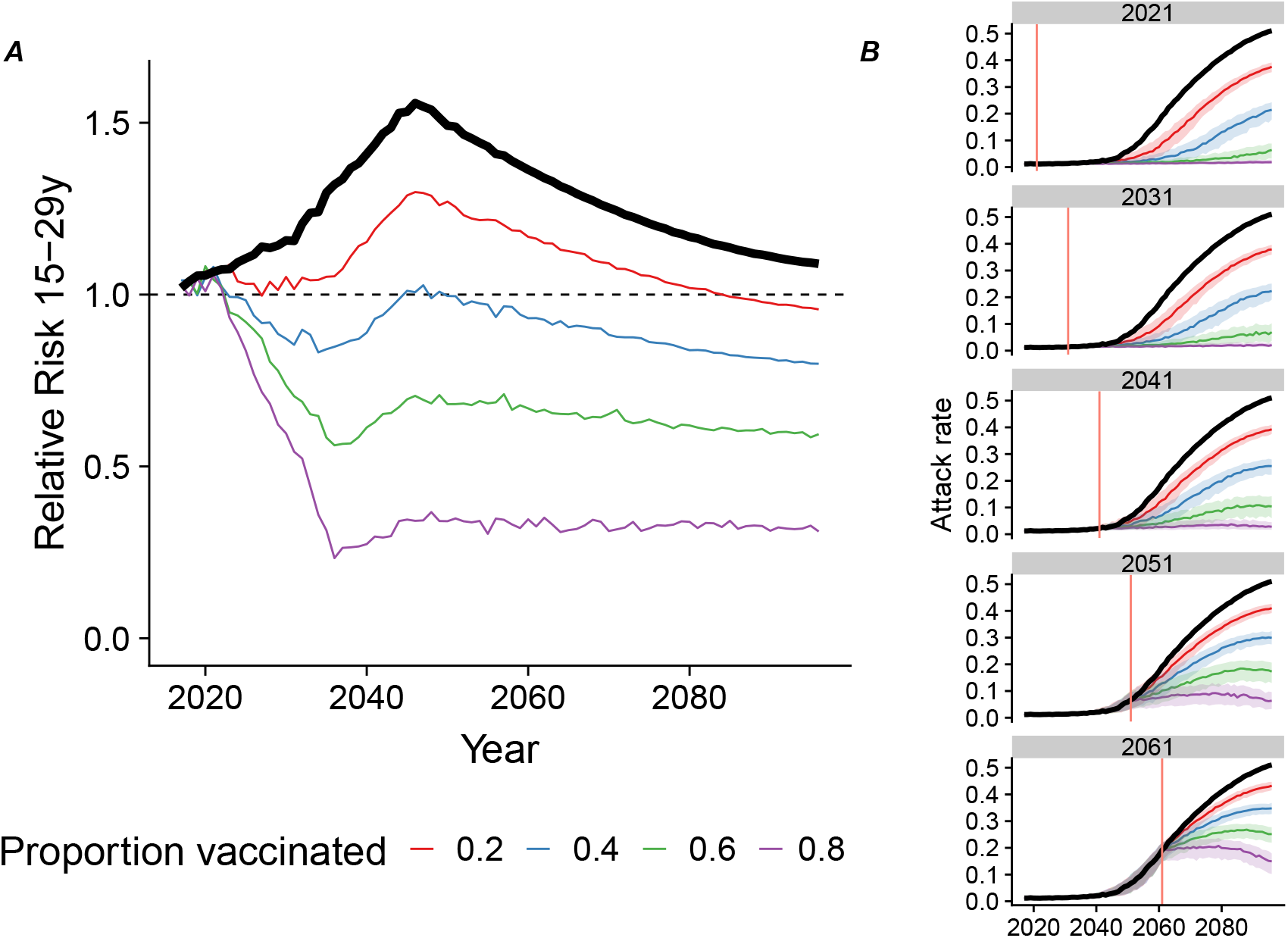
Consequences of implementing a targeted vaccination program among 15-year-old-girls from 2021 onwards with various levels of effective vaccination coverage (from 20 to 80%) compared with no vaccination (thick black line): (A) relative risk of ZIKV infection during an outbreak in the 15–29 year age group compared with the general population and (B) attack rate of ZIKV among reintroductions resulting in outbreaks (median, interquartile range, with a threshold of 1%), when vaccination is introduced from 2021, 2031, 2041, 2051 or 2061 onwards (red vertical line).

## 4 Discussion

In this mathematical modelling study, we show that a new ZIKV outbreak in Nicaragua would affect proportionally more women in the young reproductive age range (15–29 years) than the general population, owing to the age-dependent infection pattern and population renewal. The risk of a new ZIKV outbreak in Nicaragua, after reintroduction, will remain low before 2035 because of herd immunity, then rise to 50% in 2047. If protective immunity to ZIKV decays with time, ZIKV recurrence could occur sooner. Timely introduction of targeted vaccination, focusing on females aged 15 years would both reduce the risk of adverse congenital outcomes and extend herd immunity, mitigating the overall risk of an outbreak and resulting in lower attack rates if an outbreak occurs.

### 4.1 Strengths and limitations of the study

A strength of our approach is that it allows for the full propagation of uncertainty from the initial data into the risk assessment, by transferring the posterior distributions of the parameters from the deterministic model fitted to surveillance and seroprevalence data on the 2016 epidemic into the ABM used for simulations. Roche et al. showed that, when a sufficiently small time step was chosen, stochastic and deterministic models using the same parameter values led to similar results (Roche et al., 2011b). Additionally, we benefited from the availability of high quality data from population-based surveys that included participants from age 2 to 80 years in Managua, Nicaragua. The age-stratified seroprevalence data allowed us to investigate the risk in different age groups and better assess the evolution of the age-specific immunity, which is crucial when studying adverse congenital events caused by ZIKV infection during pregnancy.

We chose a simple approach based on an SIR structure, similar to the model used by Netto et al., to focus on the dynamics of infection and immunity in the human population. We did not model vector populations and behavior explicitly, as in some other studies (Kucharski et al., 2016; Champagne et al., 2016; Ferguson et al., 2016). This simplification limits the mechanistic interpretation of the epidemic parameters, but provides a phenomenological description of the transmission dynamics. We believe that this approach is appropriate because our main objective was to determine the risk of an outbreak after reintroduction of ZIKV, which is mostly influenced by the level of protective immunity in the human population. We acknowledge that the future occurrence of ZIKV in the area also depends on the presence of a competent vector. Our choice is supported by sensitivity analyses that show that more complex model structures (delayed SIR and Ross-MacDonald-type models) were not superior to a simple SIR structure in describing the 2016 ZIKV epidemic of Managua (Appendix A.6). Similarly, Pandey et al. (2013) showed that additional model complexity does not result in a better description of the dynamics of transmission of dengue virus (another Aedes-borne virus) in a human population compared with a SIR model (Pandey et al., 2013). In our model, the transmission rate (*β_a_*) captures both human-mosquito and mosquito-human transmission; we assumed a constant transmission rate, as observed in the 2016 outbreak.

Another limitation of our model is that we did not take migration or changes in population distribution into account in our model. An influx of people with lower levels of protective immunity or higher birth rates would increase the speed at which the population becomes susceptible again. Nicaragua has an urbanization rate that exceeds the world average (Maria et al., 2017). If rural populations have lower seroprevalence for ZIKV, as was shown in Suriname (Langerak et al., 2019), an inflow of rural inhabitants into Managua could increase the risk of ZIKV outbreaks. Uncertainty remains, as factors such as the political instability in Nicaragua could drive migration and influence disease transmission, as we currently observe in Venezuela and bordering countries (Tuite et al., 2018).

### 4.2 Interpretation in comparison with other studies

This study shows that the lower attack rate of ZIKV in children than in adults will hasten the emergence of a population that will be fully susceptible to infection, especially if immunity is not lifelong. The advantage of our approach is that we used the age-specific attack rates to model the processes of ageing in relation to protective immunity to ZIKV explicitly. Even with lifelong immunity, our model predicts that children aged 0–14 years will become entirely susceptible by 2031 and 15–29 year olds by 2046. In future outbreaks, the attack rate will then be highest amongst 15–29 year olds, including women who will be at risk of ZIKV infection in pregnancy. If immunity wanes, the time until the next ZIKV outbreak will be reduced and, in that case, the distribution of infection risk would be more equal across age groups (Fig. 4). Several authors have studied the time to a next ZIKV outbreak, but none studied the effect of the loss of immunity over time in relation to age. Assuming lifelong immunity, our estimates of the time until the risk increases are similar to the 12–20 years before re-emergence estimated for French Polynesia (Kucharski et al., 2016). Netto et al (2017) used an SEIR model to show that in Salvador, Brazil, the effective reproduction number was insufficient to cause a new outbreak during the “subsequent years” (Netto et al., 2017). Lourenço (2017) showed the same for the whole of Brazil: herd immunity should protect the population from a new outbreak in the coming years (Lourenço et al., 2017). Ferguson et al. (2016) concluded that the age distribution of future ZIKV outbreaks will likely differ and that a new large epidemic will be delayed for “at least a decade” (Ferguson et al., 2016).

Other ZIKV vaccination studies confirm our findings. However, they do not show the effect in risk groups nor assume herd immunity from previous outbreaks like we did here; Durham et al. (2018) showed that immunizing females aged 9 to 49 years with a 75% effective vaccine and a coverage of 90%, would reduce the incidence of prenatal infections by at least 94%. Similarly, Bartsch et al. (2018) showed that women of childbearing age or young adults would be an ideal target group for vaccination.Valega-Mackenzie and Ríos-Soto (2018) formulated a vaccination model for ZIKV transmission that included mosquito and sexual transmission. They found that vaccination works if well administered, both when sexual transmission is most important and when vector-born transmission is most important.

### 4.3 Implications and future research

Our finding that people in the 15–29 age range are more at risk of infection implies that we expect a higher number of congenital abnormalities due to ZIKV infection. Thus, vaccine development efforts should be increased. Our conclusions are drawn based on data from Managua, Nicaragua, but should be relevant to many regions in the Americas and the Pacific that have documented high post-epidemic levels of seropositivity (Aubry et al., 2017; Netto et al., 2017; Saba Villarroel et al., 2018). In regions where ZIKV has not yet caused an epidemic but competent vectors are present, vaccination would be in place as well. Further age-stratified seroprevalence studies, using sensitive and specific tests and with longitudinal follow-up, are needed to improve our understanding of ZIKV antibody distribution in populations and to quantify the duration of immunity. This information will provide important information to improve mathematical modeling of ZIKV risk.

ZIKV vaccine development faces considerable hurdles. First, the evaluation of vaccine efficacy has stalled because the reduced circulation of ZIKV has reduced the visibility of ZIKV-associated disease (Cohen, 2018). Second, it remains unclear if neutralizing antibodies induced by vaccination are sufficient to protect women against vertical transmission and congenital abnormalities (Diamond et al., 2018). Third, it is not clear whether or how vaccine-induced antibodies against ZIKV will cross-react with other flaviviruses. To move vaccine development forward, we need to find regions where disease will occur to be able to conduct trials. This requires identifying populations that are at risk, and implementing surveillance there. These can either be regions where ZIKV is endemic, or where ZIKV outbreaks are likely to occur; throughout the Americas, there might be regions that did not experience an outbreak, but do have suitable conditions such as competent vectors. Conducting vaccine trials in disease outbreaks is complex, but there are tools to facilitate planning (Bellan et al., 2019). ZIKV in an endemic setting, such as in Africa and Asia, could prove a suitable setting as well. However, ZIKV circulation in endemic setting is not well described and the occurrence of adverse outcomes in this context is less documented Counotte et al. (2018). Further research in understanding the transmission of the virus in an endemic context is therefore needed.

### 4.4 Conclusion

Preparedness is vital; the time until the next outbreak gives us to opportunity to be prepared. The next sizeable ZIKV outbreak in Nicaragua will likely not occur before 2035 but the probability of outbreaks will increase. Young women of reproductive age will be at highest risk of infection during the next ZIKV outbreak. Vaccination targeted to young women could curb the risk of a large outbreak and extend herd immunity. ZIKV vaccine development and licensure are urgent to attain the maximum benefit in reducing the population-level risk of infection and the risk of adverse congenital outcomes. The urgency of ZIKV vaccine development increases if immunity is not lifelong.

## 5 Acknowledgments

Calculations were performed on UBELIX (http://www.id.unibe.ch/hpc), the HPC cluster at the University of Bern. MJC received salary support from the Swiss National Science Foundation (project grant 320030_176233).

## 6 Competing interests

## Appendices

## A Appendix

### A.1 ABM algorithm

Here, we provide the pseudo code of the ABM (Algorithm 1).

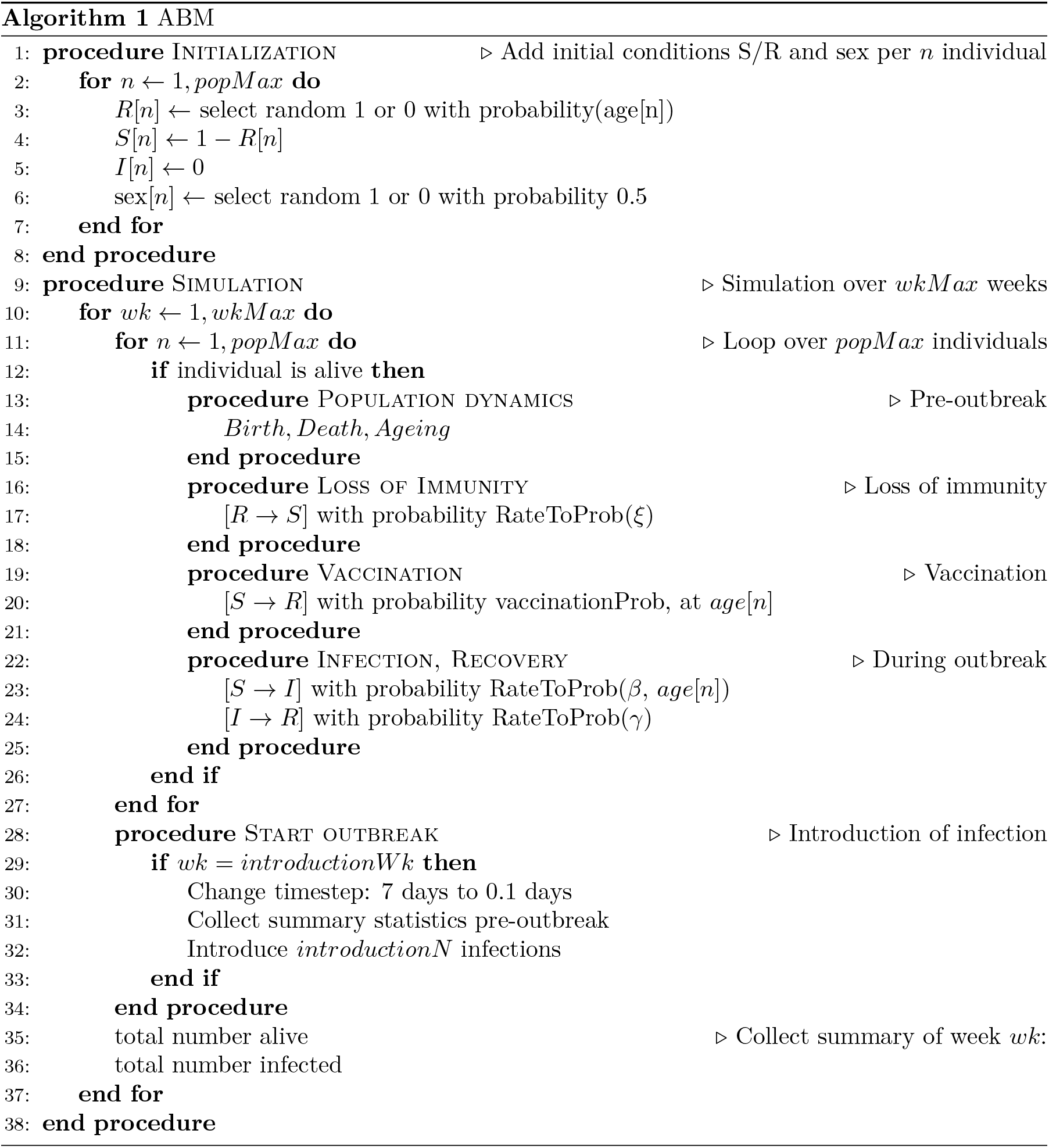

### A.2 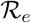

We used the next generation matrix method described by Diekmann et al. to calculate 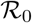 (eq. 8 – 10). *β*_1_ is the transmission rate for the 0–14 age group; *β*_2_ for the >15 group; *γ* is the common recovery rate.

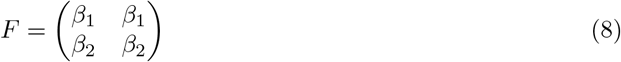

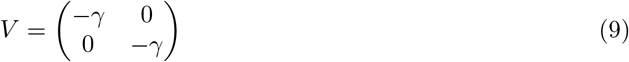

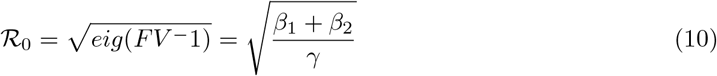

### A.3 Loss of immunity scenarios

We explored plausible scenarios of loss of immunity (Fig. A.1).

**Figure A.1:**
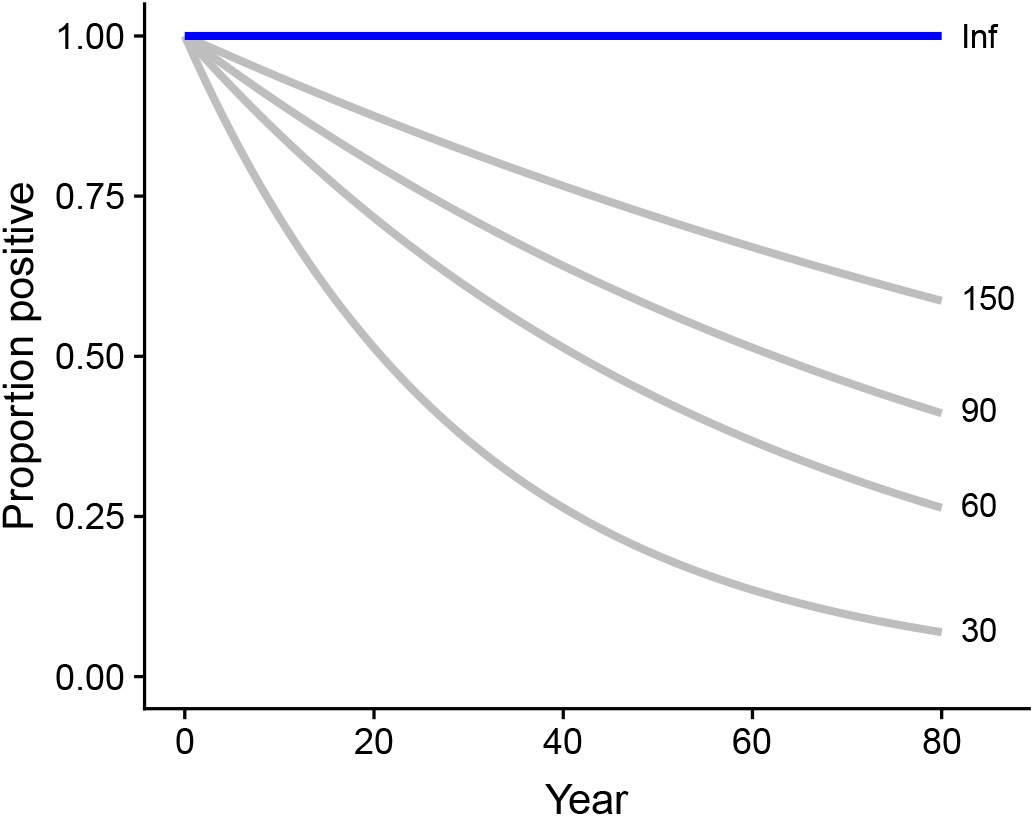
Different scenarios of loss of immunity. No loss of immunity (blue) and scenarios explored (grey, exponential function with mean durations 30, 60, 90 and 150 years).

### A.4 The number of infections introduced does influence the probability of an outbreak, but not the attack rate of successful outbreaks

The proportion of outbreaks (1% threshold) after introduction depends on the number of infections introduced; the attack rate of the successful outbreaks does not depend on the number of infections introduced (Fig. A.2).

**Figure A.2:**
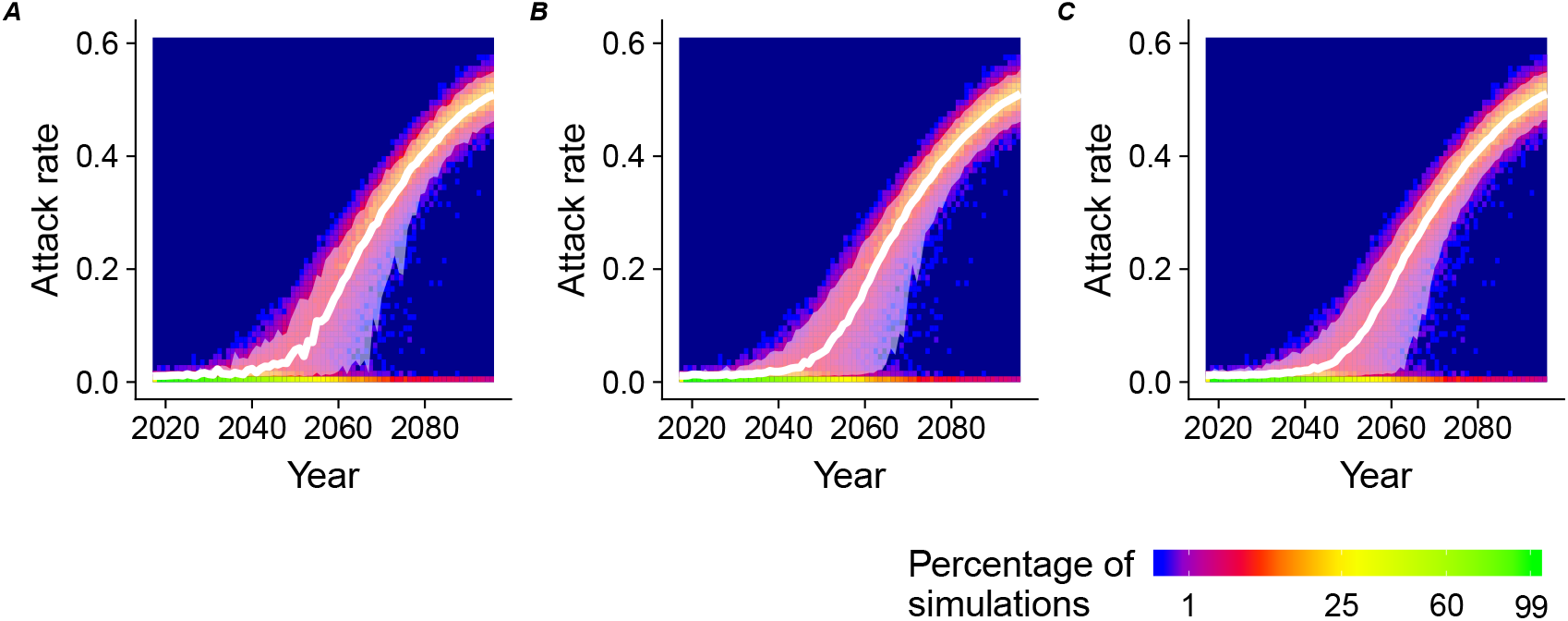
Attack rate over time for the introduction of (A) n=1, (B) n=5, (C) n=10 infections.

### A.5 Once per simulation introduction vs once per year introduction of infection

Using the ABM (model B) we explored the effect of yearly introduction of one infectious individual in the population (n=10,000). In the main text, we assumed a single introduction of n individuals per simulation. Here we introduce on a yearly basis the infectious individuals; previous outbreaks during the simulation affect the likelihood of a next outbreak and observed patterns are more stochastic. However, the pattern of the attack rate over time remains similar to the findings of the once/simulation introduction (Fig. A.3); the variation is larger due to a more stochasticity.

**Figure A.3:**
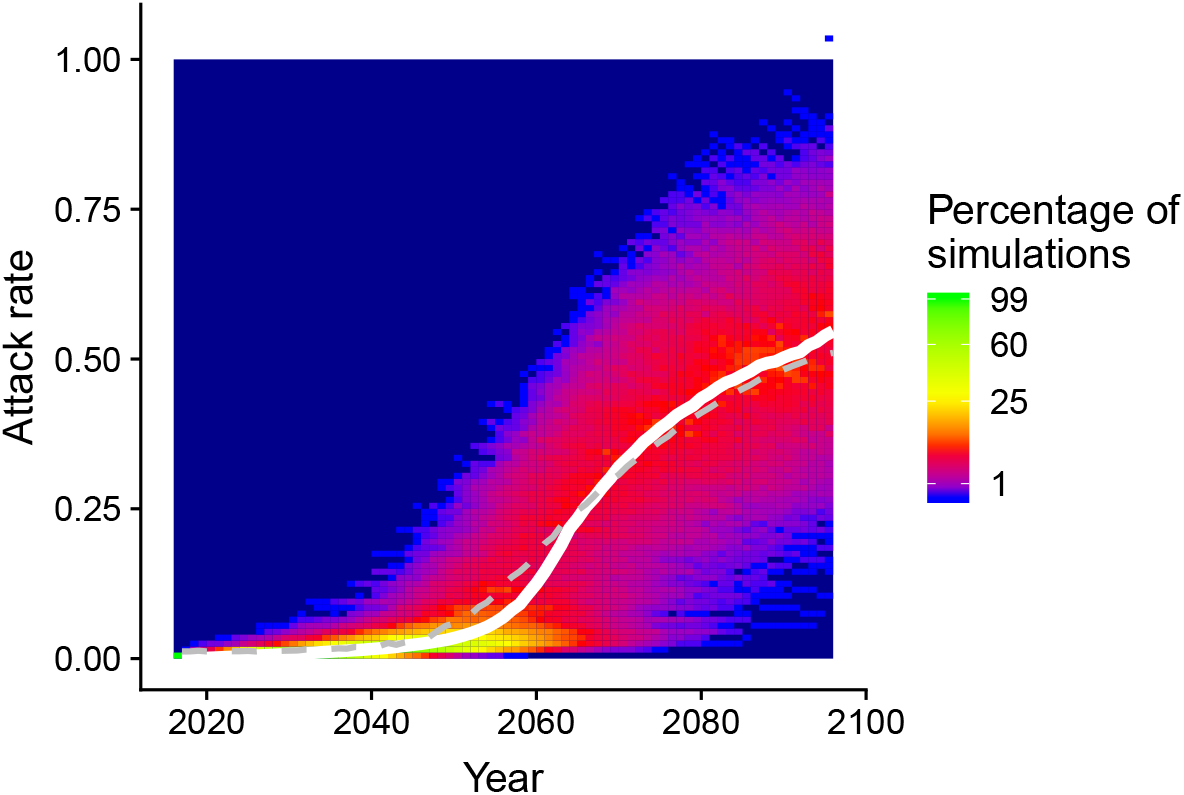
Heat map of attack rate per time for simulations where every year one infectious individual is introduced in a population of 10,000. The median (white line) of this simulation is compared with the median of the simulation where once per simulation an infection is introduced (dashed grey line).

### A.6 Comparison of SIR model with SEIR model and the Pandey model

We compared the SIR model with a SEIR and model that explicitely models the vector; the Pandey 2013 model as implemented in Champagne et al. (2016) (Champagne et al., 2016). The model fit of the more complex models does not outperform the fit of the simplest (SIR) model (Table A.1), justifying the model choice.

**Table A.1:** Model comparison

| Model | LOOIC (SE) | ΔLOOIC (SE) |
| --- | --- | --- |
| SIR | 95.2 (8.7) | Ref. |
| SEIR | 93.9 (8.9) | -1.3 (1.6) |
| Pandey | 99.5 (8.3) | 4.3 (3.6) |

